# City and forest great tits respond differently to artificial light at night: insights from a common garden study

**DOI:** 10.64898/2026.09.29.753716

**Authors:** Therese Solberg, Pepijn van der Ham, Erik Matthysen, Marcel Eens, Kamiel Spoelstra, Kees van Oers, Marcel E. Visser, Barbara M. Tomotani

## Abstract

Artificial light at night (ALAN) disrupts the natural light-dark cycle, which is the most important cue for synchronizing circadian clocks. If populations most exposed to ALAN adapt to get less disturbed by it, we expect forest and city populations to differ genetically. We tested this prediction in a common garden experiment using a songbird, the great tit (*Parus major*), originating from forest and city populations. We simulated ALAN exposure during the breeding season by keeping the birds in pairs under ALAN or dark nights for three months and then repeated the experiment in the following year with the same pairs under the opposite treatment. The advancement of daily onset under ALAN disappeared over time for city birds, but not for their forest counterparts. For these same birds, we measured their onset of daily activity under constant dim light and shift in activity in response to light pulses and showed that the city/forest difference in behavioural response was not reflected in any clock property measured. This indicates that city and forest birds differ genetically in the downstream behavioural response to ALAN, and not in the circadian clock. Thus, urbanization is unlikely to select for divergence in circadian clock properties in songbirds.

## Introduction

The global increase in artificial light at night (ALAN) over the past century has led to nights that are no longer truly dark [1,2]. Due to its impact on behaviour and physiology on a range of taxa, ALAN has gained much attention and has been recognized as an environmental stressor and a potential threat to biodiversity [3–5]. Several studies have reported profound effects of ALAN on bird physiology and behaviour [6–8], where one of the main phenotypes observed is a change in daily activity rhythms [9–11]. Interestingly, changes in daily rhythms in response to ALAN seem to differ between animals living in city and forest environments. For example, forest European blackbirds (*Turdus merula*) displayed higher nocturnal restlessness when exposed to ALAN compared to their city counterparts [7]. A similar difference is observed in city and forest great tits (*Parus major*), where an earlier onset in activity is seen in forest birds when experimentally exposed to ALAN when compared to their city counterparts [12]. This pattern could indicate an adaptation of city animals to higher levels of ALAN.

One of the proposed mechanisms that could explain how ALAN affects daily activity patterns is through an impact on the underlying circadian clock. Daily rhythms are not simply a response to the environment, but are generated through an endogenous timekeeping system, the circadian clock, that produces endogenous rhythms with a period of ∼ 24 hours in physiology and behaviour, known as circadian rhythms [13]. External environmental cues synchronize circadian rhythms and, importantly, the most prominent cue is the light/dark cycle [14,15]. This synchronization process of daily timing to the external environment depends both on properties of the circadian clock and the strength and the timing of the external cue. For example, both the endogenous clock speed (or free running period) known as *tau* as well as the moment in at which an individual is exposed to light within the activity cycle (start, middle, end) can determine the response to ALAN [15,16].

There are at least three possibilities for ALAN to impact the rhythm (See Fig. 1): 1) ALAN directly disrupts the clock by changing the moment of expression of its molecular components, thus leading to a different phase relation between the clock and the light/dark (LD) cycle which implies a change in entrainment due to direct changes in clock properties, or due to population differences in clock properties [16–19]. 2) ALAN impacts the light signal that reaches the circadian clock, leading to a different perception of photoperiod, i.e. a change in entrainment due to a change in the perceived light input (which can also be due to population differences) [16,20], in this regard it is also important to consider how and if ALAN interferes with seasonality, as daylength (photoperiod) varies across the year, diurnal birds will naturally become earlier as daylength increase in the beginning of the breeding season relative to sunrise, but later to sunrise at the end of the breeding season [21]. Hence, ALAN could interfere with the seasonal interpretation of photoperiod, e.g. a study by Dominoni et al. [22] reported seasonal variation in the impact of ALAN on the start of daily dawn song. 3) ALAN affects the activity rhythm directly, bypassing the clock, i.e. masking (direct behavioural response); this would be a plasticity of the system in response to the environmental stimuli [16,23,24].

**Fig. 1.**
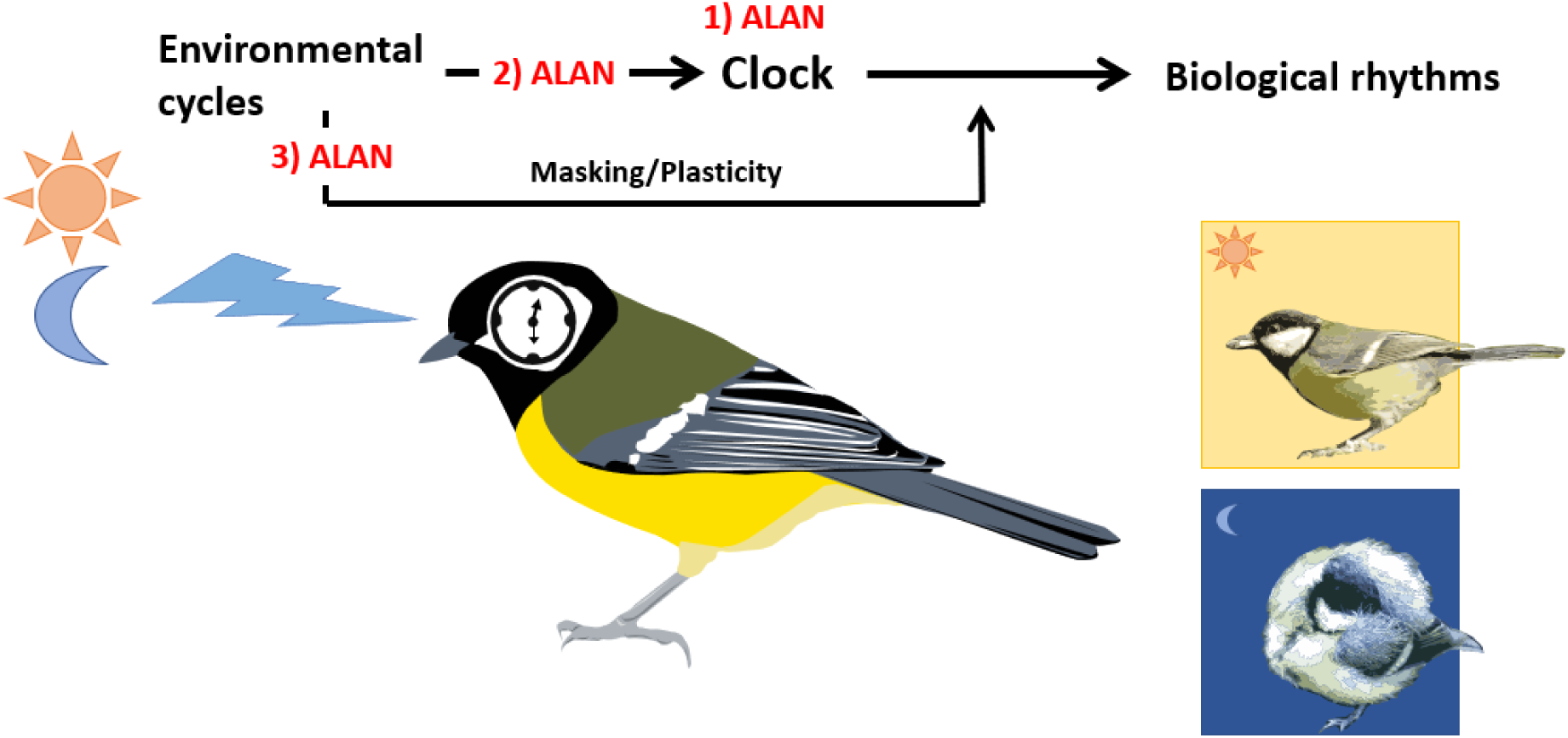
Eskinogram of circadian system where environmental LD cycle functions as input to the clock who in turn modulate output rhythms (activity/physiology) and the potential ways (scenarios) ALAN may affect output biological rhythms of activity. Where in scenario 1 ALAN affects the activity rhythm through direct effect of clock. In Scenario 2 ALAN affects the light input into the circadian system, affectively changing the perception of light, and in scenario 3 ALAN directly affects the output activity rhythm without affecting underlying clock (masking/plasticity of system). For further details see manuscript text.

It is possible to use behavioural measurements of the activity rhythm to distinguish between those scenarios, where any change in the rhythm induced by ALAN must persist in constant experimental conditions if there is a change in the underlying clock. For example, Spoelstra er al., [24] observed that upon placing birds previously exposed to ALAN in constant dim light conditions, the phase of the activity rhythm directly reverted to phase of the rhythm birds expressed even before exposed to ALAN. If the clock phase itself was changed, the advancement of the onset of activity caused by ALAN under synchronized (LD) conditions should continue after a change from LD to constant experimental conditions (change in entrainment through the clock: ALAN 1 Fig.1) or the light perception (ALAN 2: Fig.1) [16,23]. This suggests that the phase of the circadian clock itself was not affected (ALAN 3 in Fig. 1) [24].

For great tits, a recent study suggested a difference in light sensitivity between forest and city individuals [20]. It showed that when put under constant conditions, city and forest female great tits differed in their activity rhythm in the first days of the experiment, showing differences in what is known as “aftereffects of entrainment” [20]. This phenomenon reflects the history of the previous condition that the animal was exposed to, for example dark nights versus ALAN, and can be interpreted as a form of “clock memory”. In practice, when the animal is placed in constant conditions it retains a similar time of start and end of activity and a 24h period for some days. Then, it gradually reverts to the endogenous circadian period. The stronger the aftereffects, the more time it takes to revert [20]. In the study, the activity rhythm of forest birds immediately reverted to their internal clock time, no longer following 24h rhythm, while city birds remained with a 24h period for several days. The stronger aftereffects in city birds indicate a lower sensitivity to light in city birds [20].

It is widely recognized that birds alter their daily behaviour because of ALAN [9–11], but the consequences of this are not fully understood. Thus, a first step to understand if and how ALAN shapes microevolutionary processes in the city environment is to assess whether differences in responses to ALAN between city and forest populations are of genetic origin. Here we tested this through a multi-population common garden experiment. We raised great tits originating from different city (n = 5) and forest (n = 6) populations in the Netherlands and Belgium and exposed the same individuals during the breeding season to both ALAN and dark nights using a within-subject design. Birds from the same city and forest populations have already been reported to differ in other traits such as size [25], but it is yet to be determined if they genetically differ in daily timing as a consequence of ALAN. Our aim was to determine whether birds originating from city and forest populations differ in their response to ALAN when it comes to daily timing. To test whether differences are related to underlying differences in clock properties or due to a direct effect of the environment (masking), we measured three clock properties: free running period (reflecting internal clock speed), first onset of activity in constant dim light conditions (reflecting clock timing), and phase responses to a light pulse (demonstrating the clock response to light).

## Materials and methods

### Animals and common garden design

In the spring of 2021 unincubated great tit eggs from five city (Amsterdam lat: 52.372, long: 4.893, Antwerp lat: 51.16, long: 4.4, Rotterdam lat: 51.92, long: 4.48, Delft lat: 52.012, long: 4.359, Utrecht lat: 52.09, long: 5.121) and six forest populations (Bennekom lat: 52.003, long: 5.078, Boshoek lat: 51.13, long: 4.52, Hoge Veluwe lat: 52.041, long: 5.856, Liesbos lat: 51.582, long: 4.7 , Oosterhout lat: 51.873, long: 5.584, Vlieland lat: 53.295, long 5.051) were shipped to the Netherlands Institute of Ecology (NIOO-KNAW) and placed in foster nests in the nearby forest of Bennekom (the Netherlands), where the eggs were incubated and the nestlings raised by the foster parents. For more details on city and forest locations see Supplementary Material 1. On day 10 post hatching, the birds were brought to the NIOO-KNAW for hand rearing and were used in the experiment as adults in the following breeding season. More details of the common garden study are described in [25,26] and for further information about birds see Supplementary Material 2 Table S1.

### ALAN experimental design

The experiment was divided in two parts: 1) investigate if ALAN affects daily temporal activity patterns of city and forest background great tits differently across the breeding season, and 2) individual clock characterization (see Supplementary Material 2). The same experiment was conducted for two consecutive years (breeding season 2022 and 2023) using the same individual birds, but with reverse ALAN treatments (birds exposed to ALAN in 2022 were exposed to dark nights in 2023 and vice versa).

#### Part 1: daily activity patterns across the breeding season

In the experiment in 2022 we used 35 forest and 35 city birds, placed in male/female pairs (forest pairs: n=16, city pairs n=15, mixed pairs n=5). Due to shortage of birds to form pure origin pairs and to avoid pairing siblings, we had to create a few mixed pairs. Because we remained with unpaired birds, we formed 2 additional pairs using one non-common garden bird in each. The only purpose of these additional birds was to make sure all data was collected using male/female pairs and these extra birds were not included in the analysis. Some birds died during the two years (4 forest; 3 males and 1 female, 4 city: 3 males and 2 females). Hence, in 2023 to complete pairs where the partner had died, 6 additional forest birds (5 males, 1 female) where added to form pairs. Birds that died were included in the year 1 dataset when they were still healthy, and additional birds of the correct genetic background were included in the year 2 dataset (Supplementary Material 2 Table S1)

Birds were placed in male/female pairs in 36 climate-controlled aviaries (2 m x 2m x 2.25m) with one pair per aviary, at NIOO-KNAW. From early-March until mid-May birds were exposed to an artificial light-dark (LD) cycle following the natural photoperiod increase of the Netherlands and kept at a constant temperature of 13°C. The climate-controlled aviaries were equipped with 3 TL lights for daylight emitting ∼1000 lux (measured from mid-point aviary) from sunrise to sunset. The aviaries were also equipped with a light bulb that went on 30 min before sunrise and went off 30 minutes after sunset to mimic twilight. Half of the pairs (8 forest, 7 city and 3 mixed) were exposed to ALAN treatment by a nightlight emitting 1.5 lux measured from mid-point of aviary, the other half of the pairs were exposed to dark nights as control (for light spectra info see Supplementary Material Figure S4). The ALAN intensity was based on what we would expect the birds to experience in the wild [27,28]

For daily activity patterns we used feeding activity as a proxy, as it correlates with daily activity (See Supplementary Material 3). Individual feeding activity was monitored from March until mid-May. The moment of the onset of individual feeding activity was measured by using a radio frequency identifier (RFID) system (LID665 Decoder and ANTC40 antenna, DorsetID, Aalten, NL) placed on the feeder inside the aviary. Each bird was equipped with a unique transponder tag (125 kHz, 2.6mm EM4102 PIT tag, Eccel Technology, Leicester, U.K) on its leg, which was read every time the bird came in contact with the feeder – providing us with individual activity data per bird.

#### Part 2: Clock Properties

To measure if there was a difference in circadian clock properties between city and forest birds in response to ALAN, at the end of the recordings in the aviaries, birds were individually placed in a different set up. We measured their endogenous circadian rhythms in absence of time cues, for setup details see [29]. This was done for both years – having clock properties measured after control and ALAN treatments for the same individual birds. In brief: all birds were kept under dim light for 24 hours per day for 21 days. Each cage was equipped with a lamp calibrated to provide 0.5 lux at perch level to enable birds to find food and water. Feeding times followed a schedule that prevented birds to synchronize their activity to the moment they received food but remained in the time window 09:00-14:00 (CEST). Activity was continuously monitored through passive infrared sensors (PIR), detecting infra-red radiation in their field of view (entire cage) to gather activity data. In order to test the sensitivity of the circadian clock to light, 11 days after part 2 started a light pulse of 4h duration, 800 lux was given at circadian time 0 (CT0), which is the predicted time of activity onset if there was no light pulse. Since all birds had a clear window of activity and rest, it was possible to predict CT0 using the onsets of activity and the bird circadian period prior to the light pulse. Here, the expectation was that more sensitive clocks would produce a larger shift in the phase of the onset of activity [15,30].

Because the number of birds was larger than the number of cages available in the set-up, birds had to be measured in two groups (Group A and Group B), where Group A was moved into the setup on the 1st of June and group B was moved into the setup on June 29th. To minimize differences between the measurements of these two groups, habitat (city or forest and genetic population were divided as evenly across groups as possible, and we kept Group B in the climate-controlled aviary with the photoperiod fixed on the 1^st^ of June, to prevent any effect of daylength on clock properties [31]. The bird groups stayed the same across years (e.g. individuals in Group A were the same in 2022 and 2023). Due to technical problems, we lost the first three days of recording for group A in the 2023 cohort.

#### Data analysis

All statistical analyses were conducted in R (version 4.3.0). For analysis of statistical significance, we used linear mixed effect models with the R package “lme4” using “lmer” function. P-values were obtained through Kenward Roger approximation, where models were compared by dropping terms of interests through backwards selection in a stepwise manner – starting with full model using the R package “pbkrtest” with the “KRmodcomp” function [32]. Assumptions for linear model were tested for all models using qqplots and histograms of residual values, and although there were some deviations, linear mixed models through lme4 package and Kenward roger approximation have been shown to be robust against them [33,34].

#### Daily activity – onsets and offsets

We extracted the first and last RFID recording per bird per day and converted them to decimal hours (hereby referred to as onset and offset). We ran two separate linear mixed effect models, one for onset and one for offset. Each full model contained the fixed effects Treatment (control or ALAN), Habitat (city or forest), days since the 31^st^ of March, herby referred to as Aprildate (to test for seasonal changes), year (2022 or 2023) and sex (male or female). To test if ALAN affected daily activity patterns differently throughout the breeding season and across habitat of origin – we included the 3-way interaction between Treatment, Habitat and Aprildate and the two-way interactions of all combinations with these three terms. In order to account for effects of multiple measurements of the same individual (average recording per individual per year: 48 onsets, ranging from10 to 54 measurements, 48 offsets ranging from 11 to 54) as well as for the fact that we measured two birds in the same aviary, we included individual ID nested within aviary as a random effect. Likewise, to account for family and area of origin effects, we included the genetic mother ID nested within the population of origin (the specific city or forest area the egg came from). For significant interactions *post hoc* analyses were conducted through pairwise comparisons of estimated marginal trends.

#### Clock properties

The clock properties were calculated using the software Chronoshop 1.1 [35]. We investigated three clock properties: 1) the first onset of activity in constant conditions – reflecting the timing of the internal clock in relation to predicted sunrise (the moment of sunrise at which the lights in the aviaries would have switched on), as when put in constant conditions the phase (timing) of clock during preceding LD conditions will be revealed [16]. 2) free running period or *tau* (the “speed” of the clock) and 3) phase shift in response to 4-hour light pulse at CT0, where individuals that show larger shifts in when they first become active in response to the light pulse (i.e. larger phase shifts) have biological clocks that are more sensitive to external stimuli. For calculating activity onset, the software calculates the centre of gravity per cycle positioned at the mean vector angle. Then it estimates the activity onset by going 0.5 cycles back in time detecting the moment the activity counts (smoothened by a running mean adjusted for best fit) surpasses the average activity of the current cycle. Free-running period was obtained by excluding first days after LD with aftereffects under standard experimental conditions (when activity rhythm might be influenced by previous experimental conditions) and then calculated using the Sokolove and Bushell (S&B) algorithm [36] – only data prior to the light pulse was used. The software calculates phase shift by “stacking” onsets over multiple cycles and calculates the ‘average’ onset before and after the light pulse, where the difference between the two phases constitutes the phase shift [see 27].

We used three linear mixed effect models, one for each of the clock properties. We included the interaction between Treatment and Habitat (to test if there was a difference in the effect of ALAN between city and forest birds), as well as Year (2022 and 2023), group (A or B) and Sex (male or female) as covariates. Like in the analysis of rhythms in feeding activity, effects of repeated measures of the same individuals over two years as well as family and area of origin effect were accounted for by including individual ID nested within Mother ID, nested within area of origin (specific city or forest the egg came from) as a random effect.

#### Individual onset sensitivity to ALAN and its relation to clock properties

Next, we wanted to test if ALAN sensitivity (plasticity) phenotype could be related to the clock. I.e. if a stronger individual response to ALAN would be associated with one of the clock properties. Thus, we tested whether the individual plasticity could be linked to the clock properties measured (first onset of activity, *tau*, phase shift in response to light pulse). Since we found no significant effect of either treatment or origin in any clock property, we calculated the individual mean (the experiment was done twice, in 2022 and 2023, but with reverse treatments) for each clock property. Plasticity values for each individual were obtained by fitting a linear model with onset as response and the interaction Individual ID: Aprildate – in one model per year (only using individuals present both years; See Supplementary Material 7 for individual slopes). Then we calculated the Δ onset slope (plasticity response) by subtracting the individual ALAN slope from corresponding individual control slope. Then to test if individual plasticity could be linked to clock properties - a linear mixed model was fitted with individual plasticity as a response variable and the first onset of activity, tau and phase shift as explanatory variables. In all models, the biological mother nested within area (populational origin) was included as random effect.

## Results

### Part 1: Daily activity across the breeding season

Birds with city and forest background differed in their timing of daily activity onset in response to ALAN throughout the breeding season (significant 3-way interaction between Treatment, Habitat and Aprildate, F _1, 6494.1_ = 5.21, *p* = 0.02; Fig 2A). A post hoc analysis confirmed a significant difference in slope between city and forest birds exposed to ALAN (t=-3.72, *p*= 0.0002), where city birds advanced their onset less (slope estimate= -0.019, SE= 0.001) compared to forest birds (slope estimate -0.24, SE=0.001). Under the dark night control Treatment, there was no difference in slope between city and forest birds (t=-0.36, *p*=0.72) where birds largely seemed to follow the time that TL lights switched on (day light lamps) on (Fig. 2A).

**Fig. 2:**
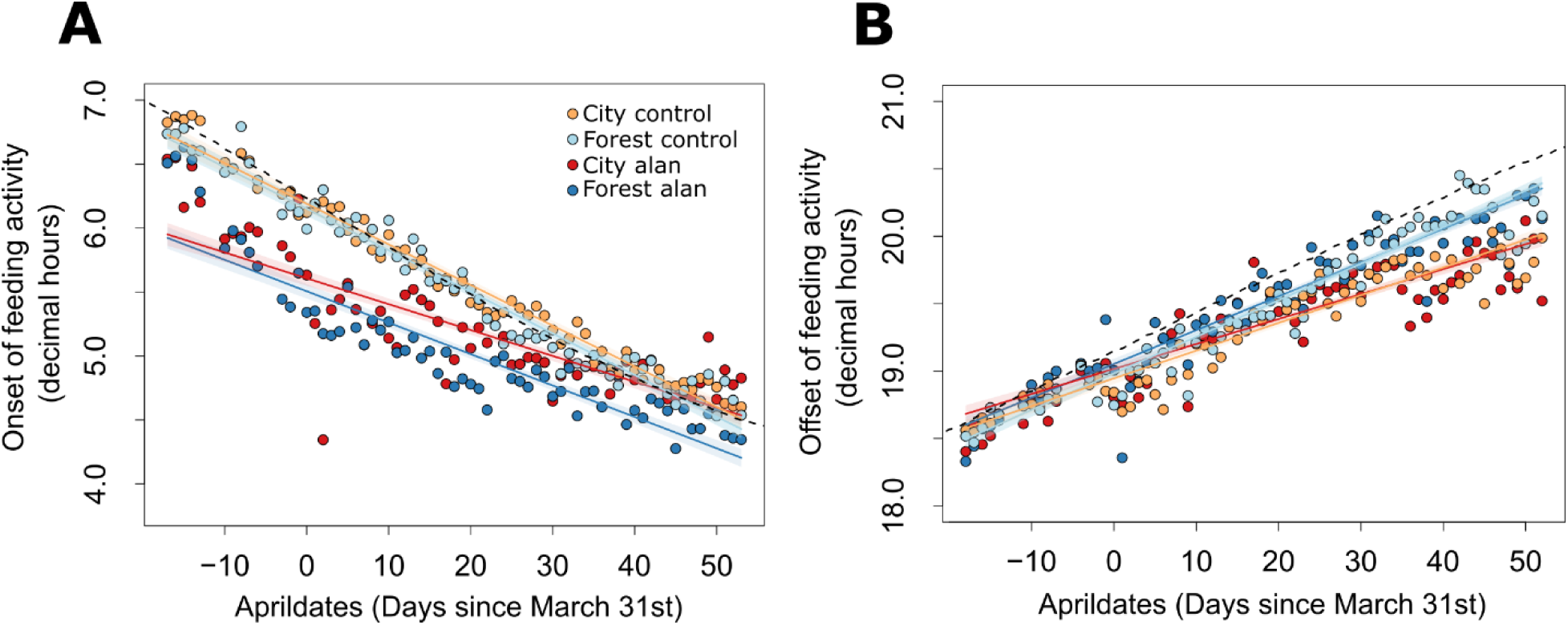
Daily raw means and regression lines for feeding activity onset (A), offset (B) with absolute time in decimal hours, across two consecutive years (2022 and 2023; with same birds reverse treatments). The dotted lines represent the photoperiod in the aviaries (mimicking the natural photoperiod in the Netherlands). Yellow represents city control, light blue represents forest control, red represents city ALAN, and blue represents forest ALAN, with regression lines corresponding to the full model predictions in the same colours.

There was no difference in mean offset between city and forest birds in response to Treatment (no significant interaction Treatment and Habitat, F_1, 6321.31_, = 0.52, p= 0.47, Fig. 2B). City and forest birds had a significantly different offset as season progressed (significant interaction between Habitat and days since Aprildate, F_1, 6488.7_ = 80.1, p= <0.0001, Fig.2B), where a post hoc analysis showed that forest birds had a later offset as the breeding season progressed (slope estimate = 0.03, SE = 0.001) compared to city birds, which ended their activity earlier (slope estimate 0.023, SE= 0.001). Finally, birds exposed to ALAN had a later offset in the beginning of the breeding season, that weakened over time when compared to birds in dark controls (significant Treatment and Aprildate interaction, F_1,6429.1_= 11.16, p= 0.0008, Fig. 2B).

For complete statistical output of the models see Supplementary Material Table 2.

### Part 2: Clock Properties

Upon placement in constant dim light conditions, city and forest birds showed similar timing in their first onset of activity in in relation to previous Treatment (no significant interaction between Treatment and Habitat, F_1, 56.3_ = 0.48, *p=* 0.49, Fig.3A) and previous Treatment did not affect the first onset of activity (Treatment, F_1,58.33_ = 0.55, *p* = 0.46, Fig.3A). There was also no difference in first onset of activity between city and forest birds (Habitat, F_1,7.28_ = 0.0001, *p*=0.99).

**Fig. 3:**
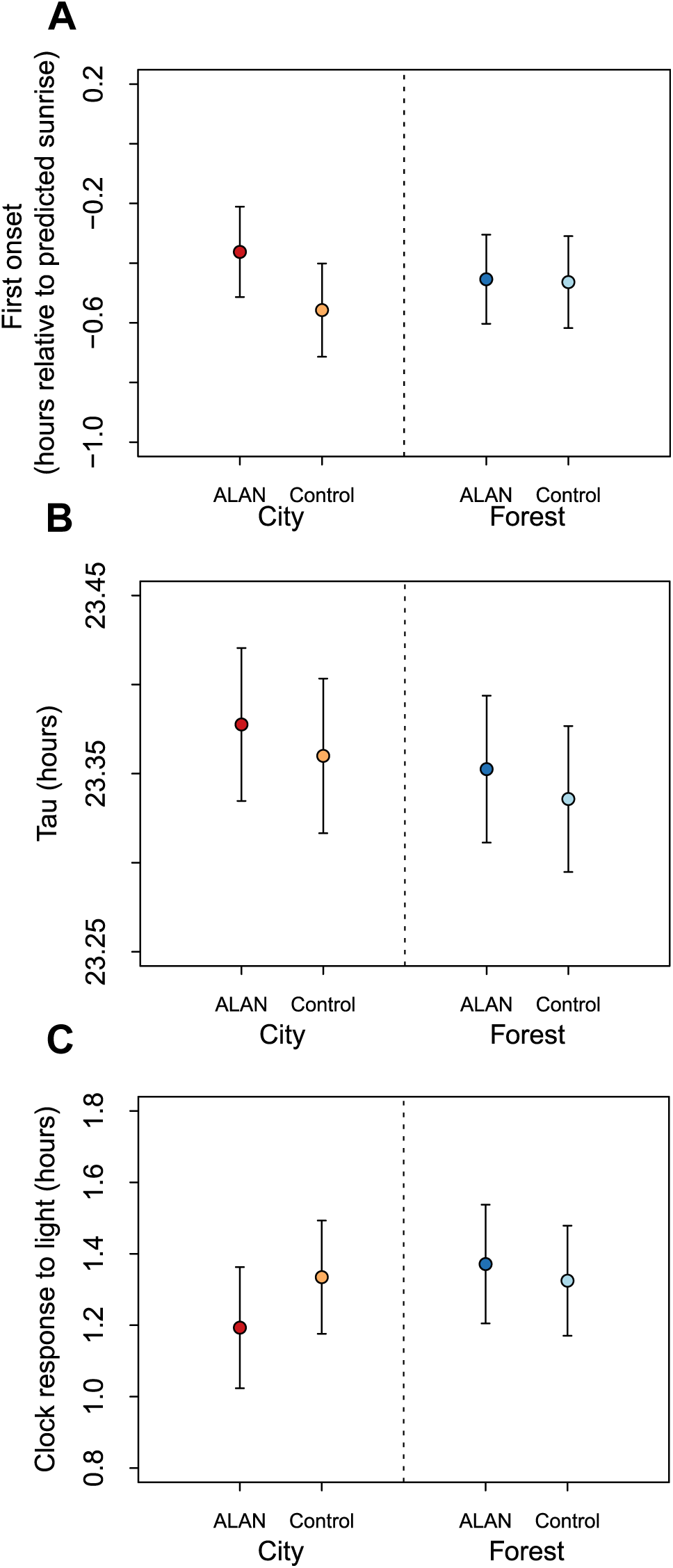
Clock properties of A) First onset relative to predicted sunrise. B) Free running period (*tau*) and C) Phase shift in response to light pulse given at circadian time (CT 0).

We found no difference in free-running period (clock speed) between city and forest birds in relation to preceding Treatment (no significant interaction between Treatment and Habitat, F_1,61.78_ = 0.0002, *p*=0.99). ALAN Treatment had no effect on free-running period (Treatment, F_1,62.91_ = 0.33, *p*=0.46), neither did we observe a difference in free-running period between city and forest birds (Habitat, F_1.7.78_ = 0.22, *p* = 0.64, Fig.3B).

There was no difference in the effect of Treatment between city and forest birds on the phase shift response to the light pulse given at CT 0 (no significant Habitat and Treatment interaction, F_1,53.16_ = 0.46, *p* = 0.5, Fig.3C). There was also no difference in phase shifts between ALAN and control Treatments (Treatment, F_1,54.1_ = 0.1, *p* = 0.75, Fig.3C), or city and forest birds (Habitat, F_1,7.34_ = 0.18, *p*= 0.68).

For complete statistical output of the models see Supplementary Material Table 3.

#### Part 3: Individual onset sensitivity to ALAN and its relation to clock properties

There was no significant relationship between individual plasticity (Δ onset slope) in response to ALAN and the clock properties analysed (first onset of activity, *tau* or phase shift response to light pulse) (See Table 1)

**Table 1:** Statistical output for the plasticity to ALAN in relation to clock properties (First onset of activity, phase shift and *tau*): Statistics are given at the point of exclusion of the model, using Kenward Roger approximation through the function “KRmodcomp”. Biological mother nested within area (population origin) are included in each model as random effects. None of the clock properties measured could be linked to the ALAN sensitivity phenotype.

| **$\Delta$ onset slope** | **Ndf** | **ddf** | **F-value** | **p-value** |
| --- | --- | --- | --- | --- |
| First onset of activity | 1 | 19.06 | 0.72 | 0.41 |
| *tau* | 1 | 38.76 | 0.03 | 0.84 |
| Phase shift | 1 | 7.74 | 2.52 | 0.15 |

## Discussion

In this common garden study, we found no differences in light sensitivity of the circadian system between city and forest birds, but forest birds had a significantly stronger response to the presence of low ALAN conditions over the course of the breeding season relative to city birds. Independently of treatment, city birds also ended their daily activity earlier over the course of the breeding season. None of the observed differences was associated with any of the clock properties measured suggesting that clock properties were not affected by the treatments, and that birds from the different habitat origins did not have genetically different clocks. We also tested whether individual onset plasticity could be linked to individual clock properties (See Table 1) – but individual plasticity was not associated with any corresponding individual clock properties measured.

City birds were less affected by ALAN compared to their forest conspecifics. This pattern has also been explored in other studies: Ulgezen et al. [12] reported that white ALAN had a larger effect on onset of activity for forest wild-caught male great tits causing earlier onsets of activity than observed in city conspecifics. In contrast, McGlade et al. [10] did not observe differences in onset of activity between female city and forest great tits experimentally exposed to ALAN in the field. When other phenotypes are examined, some studies have also reported city birds to be more tolerant to light at night than forest birds. In that same study, McGlade et al. [10] reported city birds to have less nocturnal restlessness in incubation behaviour. Another example of attenuated responses to ALAN is in levels of feather corticosterone, which is corticosterone (stress related hormone) which is integrated over time in feathers. Dominoni et al. [38] found that city blue tit (*Cyanistes caeruleus*) nestlings had a higher concentration of feather corticosterone than forest conspecifics when exposed to ALAN, where this higher concentration correlated with higher fledgling success. Hutton et al. [39] observed that wild caught rural house finches (*Haemorhous mexicanus*) experimentally exposed to ALAN, in a laboratory setting suppressed feather corticosterone in contrast to finches in cities – suggesting partly a developed resistance to ALAN [38,39].

By the end of the breeding season, city birds under ALAN ended up having a similar onset of activity as dark controls, while forest birds under ALAN were still much earlier, despite the fact that, at the start of the experiment, both origins under ALAN had a similar onset of activity (Fig.2A). While we are not able to test the mechanism behind this change, we can speculate on the causes. One possibility is that city birds get desensitized to the stimuli faster than forest birds. In this regard, when onsets are already advancing due to increasing daylength [21], birds need to be sensitive to ALAN to further advance their onset. Hence, the longer day length might ‘overpower’ the effect of ALAN for the less sensitive city birds. Another possibility is that ALAN interferes with the seasonal interpretation of photoperiod, where birds with a city genetic background act as if it is later in the season. Indeed, Dominoni et al. [22] reported seasonal variation in the impact of ALAN on the start of daily dawn song, with the effect of ALAN being reduced later in season. This suggests that ALAN alters the perception of photoperiod, where the shift in activity onset could potentially be related to breeding stage.

In our study we did not observe an effect of ALAN on the offset of activity (Fig.2B). In both laboratory and wild birds, light is reported to have a stronger effect on timing of onset than on offset of activity, with a higher variation in offset relative to light conditions [6,11,29]. De Jong et al. [6] proposed that the termination of activity is less influenced by light than by other factors, while Kempenaers et al. [40] observed that late singers, such as the great tit, have a higher threshold in light sensitivity compared to early singers (E.g. robin – *Erithacus rubecula*). This “threshold” hypothesis state that species such as the great tit that naturally stop singing before sunset (at relatively high levels of natural light) will not respond to ALAN as long as the artificial light intensity is lower than the natural light threshold [11]. Indeed, we observed in our study that while both city and forest birds do end their activity later as season progresses, it is never after the time that lights go off (i.e. their “sunset”) (Fig.2B). This supports the notion that if light intensity is lower than the threshold in the natural light levels that commonly offsets activity, it will not induce a response. Interestingly in our study city birds also ended their (feeding) activity earlier than forest birds as daylength increased (Fig. 2B). Again, this could be related to a seasonal effect in different perception of photoperiod making city birds perceive the ALAN treatment as if it is later in the season.

The light-dark cycle (LD) is the most prominent time cue for synchronizing the circadian clock to the external environment, a phenomenon called entrainment [15,16,41]. Entrainment depends on the organisms’ clock properties (e.g. internal clock speed, *tau*), the sensitivity or responsiveness of the system to the light cue, and the strength of the light stimulus (e.g. intensity, spectral composition, duration) [16,17,42]. Therefore, it can be expected that any change in the response to ALAN is caused by differences in circadian clock properties. However, for the parameters we measured in birds in our experiment, this was not the case (Fig.3 and Table 1). Thus, if we go back to our three scenarios in Figure 1, scenarios 1 or 2 are not what explains the difference in activity rhythms.

It remains unclear whether the impact of ALAN on onset is caused by a shift in the circadian clock or just the output rhythm [18,43]. If the different responses of our city and forest birds were caused by scenario 1, through clock effects (See Fig.1), we should see differences in either *tau* or the response to the light pulse between treatments and/or habitat origins. In birds, *tau* has shown to be related to daily timing – with shorter *tau* for earlier onset [19,20]. While the magnitude of the circadian response of an organism to a single light pulse reflects the clock’s responsiveness to entrainment stimuli [15,44]. In the case of scenario 2, there would be a difference in the first onset of activity in constant conditions, because this measurement reflects how the internal rhythm is aligned to the LD cycle [16]. The fact that no difference was observed in the measured clock properties points to scenario 3, i.e., that ALAN is affecting the rhythm directly (masking / plasticity) [23,24].

Molecular analyses of great tits experimentally exposed to different light intensities of ALAN under entrained condition revealed a shift in the daily phase of expression of the core clock gene BMAL1 but not in CRY, the other clock gene measured [18]. Similar results have also been reported in wild house sparrows (*Passer domesticus*) residing in areas with different ALAN intensity (Coker et al., 2025). In both cases it is implied that the earlier onset might be a consequence of the shift in the clock gene BMAl1. This shift is usually interpreted as scenario 1 where ALAN affects the clock directly. However, it could also be due to the effect described under scenario 2: Since the phase of entrainment can depend on season (photoperiod) [21,31] if birds perceive a longer photoperiod in the presence of ALAN (which causes their system to ‘think’ it is later in season), the shift in clock genes measured could be caused by samples being taken at different internal clock phases. In contrast, our results are more in line with scenario 3, similar to those of Spoelstra et al. [17], where forest great tits with substantially advanced onsets by ALAN exposure immediately reverted to their normal onset times (without ALAN) when placed in constant conditions. We also tested if the individual onset plasticity to ALAN treatment (Δ onset slope in climate-controlled aviaries) could be linked to the corresponding individual clock properties (see Supplementary Material 6), but no association was found. This further supports the conclusion that differences in the onset of activity between city and forest birds is not a consequence of a shift in the clock. We do however recognise the limitation of the method of using Δ onset slope, as the SE from one analysis is not carried over to the next. But our conclusions are supported by more than one result.

One important aspect to mention, however, is that the expression of the activity phenotype is likely under circadian control [24]. This can be implied because there is a limit to the advancement of the onset of activity under ALAN conditions. If there was no circadian control and ALAN affects nightly activity by simply increasing it, this would be distributed over the whole night, which is not the case. Hence, we are most likely observing effects in mechanisms downstream of the circadian clock.

Because we used a common garden design, the observed difference in activity onset between forest and city in response to ALAN suggests that this variation has a genetic basis. We explored whether a specific population could be driving the “city-forest” effect observed, but this was not the case (Supplementary Figure S4). This is intriguing, as it suggests that the great tit forest and city populations are sufficiently separated for (micro)evolutionary processes to take place (see also [21]). Despite the mobility of great tits during natal dispersal, and gene flow between city and forest great tit populations has been shown, local adaptation can still arise when gene flow is only moderately reduced along urbanized gradients [45]. Another possibility could also involve “phenotypic matching”, which involves non-random geneflow – where individuals with different genotypes choose their corresponding optimal habitat [46]. Hence, birds that are less sensitive to ALAN could be more prone to disperse into city habitats compared to more sensitive birds.

On the other hand, it is important to highlight that while our common garden study points towards heritable differences in city and forest birds’ responses to ALAN, we cannot fully discard possible maternal effects [47]. In our study we used the F1 generation of birds, where eggs were laid in their respective local environment (different egg environments). Hence, the results obtained could have been affected by maternal effects the environment where the egg was laid [41].

Finally, one important important point is whether the reduced sensitivity to ALAN in city birds is actually adaptive. While our study was not designed to test this, it is useful to speculate. Coker et al. [43] reported that birds residing in areas with high ALAN intensity had lower fitness, through lower parental provisioning to chicks and lower chick weight. If a reduced sensitivity to ALAN would mean that the parents provide more food to their young or be less inhibited in other behaviours, then a lower ALAN sensitivity could be an adaptive trait in cities. One way to test the fitness benefits of sensitivity to light would be through reciprocal transplant experiments where the fitness of local individuals is compared to transplanted individuals [47] and this would be an interesting, although methodologically challenging follow-up of this study.

To conclude, we showed that city birds are genetically less sensitive in their onset of activity in response to ALAN when compared to their forest counterparts, but this difference in response was not associated with any difference in clock properties measured. ALAN in cities might select birds to respond differently to light, however, our study does not support the hypothesis that urbanization selects for differences in circadian clock properties in songbirds.

## Authors’ contributions

TS: Formal analysis, Investigation, Writing – original draft, Writing – review & editing; PvH: Formal analysis, Investigation, Writing – review & editing; MEV: Conceptualization, Funding acquisition, Investigation, Methodology, Writing – review & editing; KvO: Conceptualization, Funding acquisition, Methodology, Writing – review & editing; KS: Methodology, Writing – review & editing; EM: Investigation, Writing – review & editing;ME: Investigation, Writing – review & editing; BMT: Conceptualization, Formal analysis, Funding acquisition, Investigation, Methodology, Supervision, Writing – original draft, Writing – review & editing

## Ethics

All experimental procedures in the lab and in the field were carried out under licenses of the Central Authority for Scientific Procedures on Animals (Project AVD 80100 2019 9005) of the Netherlands, and the Animal Welfare Body (IvD) of the Royal Netherlands Academy of Arts and Sciences (KNAW; protocol NIOO21.04).

## Declaration of AI use

AI was not used in creating this article.

## Acknowledgements

We thank Huib van Haar, Anne Walboom, Ojaswi Sumbh and João Klinke Felix for their help with the common garden study. Anne Dijkzeul, Janina Harms, Maaike van den Born, Nina Teeuw, Ruben de Wit and Vivian van ’t Westende for taking care of our birds and the technical department of the Netherlands Institute of Ecology for constructing and maintaining the experimental set-up for measuring avian circadian rhythms and all the assistance with the climate aviaries.

