## Supplementary material for "City and forest great tits respond differently to artificial light at night: insights from a common garden study": Supplemetal Material

1) **Locations of populations of origin for common garden birds**

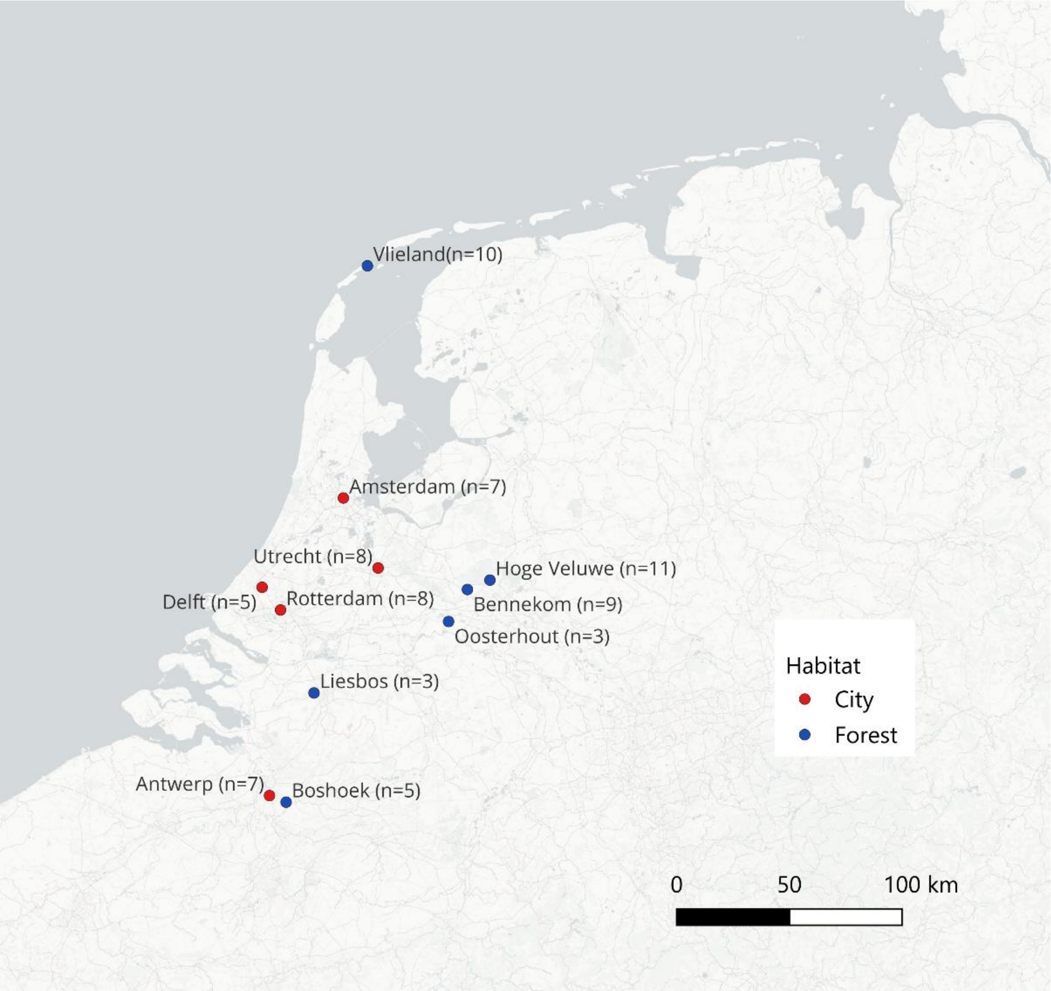

**Figure S1:** Map containing the city (red) and forest (blue) locations the eggs of the common birds originated from with sample size of birds included in study indicated. Forest sites were already existing long-term study sites that monitor great tit populations. In the Netherlands cities were all located in the Randstad region, where more than half of the country population lives, with a population of over 8 million people, and including three of the largest cities in the country. Likewise, Antwerp is the largest municipality in Belgium.

### 2) Experimental design and bird information

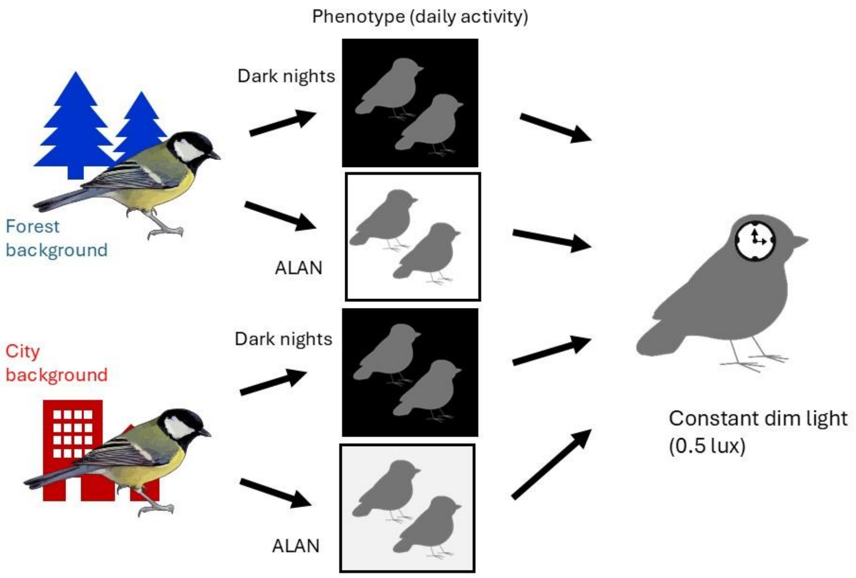

**Figure S2:** Study design of experiment. Breeding pairs of genetically city and forest great tits were formed and exposed to either dark night or ALAN. Then at the end of the breeding season, birds were put in individual cages for measurements of clock properties (First onset, *tau* and light pulse given at CT0). The experiment was conducted over two consecutive years (2022 and 2023) with the same individuals in reverse treatments – so the same individuals were exposed to both treatment conditions.

**Table S1:** Detailed Information about individual experimental birds including sex, country of origin, habitat of origin, Location of origin (population), what date the eggs where collected, family (nest), which aviary birds were kept for feeding activity recording, treatment given year 1 and 2 and info about which birds died and were replaced.

| ID | Sex | Country | Habitat | Location | EggCollection Date | Family | Pair | Treatment Year1 | Treatment Year2 | Note |
|---|---|---|---|---|---|---|---|---|---|---|
| 76819 | f | BE | Forest | Boshoek | 12042021 | 75904 | 6 | control | alan | |
| 76820 | m | BE | Forest | Boshoek | 12042021 | 75909 | 6 | control | | died after 1st year |
| 76822 | f | BE | Forest | Boshoek | 12042021 | 75909 | 11 | alan | control | |
| 76818 | f | BE | Forest | Boshoek | 12042021 | 75909 | 28 | alan | control | |
| 76821 | f | BE | Forest | Boshoek | 12042021 | 75909 | 31 | control | alan | |
| 76774 | f | BE | Urban | Antwerp | 9042021 | 75871 | 17 | alan | control | |
| 76782 | m | BE | Urban | Antwerp | 9042021 | 75872 | 18 | control | alan | |
| 76785 | m | BE | Urban | Antwerp | 9042021 | 75873 | 17 | alan | control | |
| 76784 | f | BE | Urban | Antwerp | 9042021 | 75873 | 18 | control | alan | |
| 76773 | f | BE | Urban | Antwerp | 9042021 | 75873 | 19 | control | alan | |
| 76772 | f | BE | Urban | Antwerp | 9042021 | 75873 | 22 | alan | control | |
| 76771 | f | BE | Urban | Antwerp | 9042021 | 75873 | 36 | alan | control | |
| 76777 | m | FR | Urban | Montpellier | 7042021 | 75863 | 33 | control | alan | |
| 76770 | f | NL | Forest | Bennekom | 9042021 | 75831 | 14 | control | | died after 1st year |
| 76767 | m | NL | Forest | Bennekom | 9042021 | 75831 | 20 | alan | control | |
| 76766 | f | NL | Forest | Bennekom | 9042021 | 75831 | 24 | alan | control | |
| 76776 | m | NL | Forest | Bennekom | 9042021 | 75840 | 14 | control | | died after 1st year |
| 76778 | f | NL | Forest | Bennekom | 9042021 | 75840 | 20 | alan | control | |
| 76833 | m | NL | Forest | Bennekom | 14042021 | 75973 | 19 | control | alan | |
| 76831 | m | NL | Forest | Bennekom | 14042021 | 75973 | 24 | alan | control | |
| 76834 | m | NL | Forest | Bennekom | 14042021 | 75973 | 35 | control | alan | |
| 76835 | m | NL | Forest | Bennekom | 14042021 | 75973 | 36 | alan | control | |
| 76867 | m | NL | Forest | Hoge Veluwe | 30042021 | 76396 | 6 | | alan | replaced BD76820 |

| | | | | | | | | | | |
|---|---|---|---|---|---|---|---|---|---|---|
| 76874 | f | NL | Forest | Hoge Veluwe | 30042021 | 76396 | 15 | alan | control | |
| 76871 | f | NL | Forest | Hoge Veluwe | 30042021 | 76396 | 27 | control | alan | |
| 76882 | m | NL | Forest | Hoge Veluwe | 30042021 | 76399 | 15 | alan | control | |
| 76889 | m | NL | Forest | Hoge Veluwe | 30042021 | 76399 | 27 | control | alan | |
| 76886 | m | NL | Forest | Hoge Veluwe | 30042021 | 76412 | 22 | | control | replaced BD76828 |
| 76865 | f | NL | Forest | Hoge Veluwe | 30042021 | 76435 | 10 | control | alan | |
| 76868 | m | NL | Forest | Hoge Veluwe | 30042021 | 76435 | 28 | alan | control | |
| 76870 | m | NL | Forest | Hoge Veluwe | 30042021 | 76435 | 32 | alan | control | |
| 76878 | m | NL | Forest | Hoge Veluwe | 30042021 | 76483 | 10 | control | alan | |
| 76812 | m | NL | Forest | Hoge Veluwe | 30042021 | 76483 | 26 | | control | replaced BD47429 |
| 76879 | f | NL | Forest | Hoge Veluwe | 30042021 | 76648 | 32 | alan | control | |
| 76852 | m | NL | Forest | Liesbos | 23042021 | 76717 | 3 | alan | control | |
| 76827 | f | NL | Forest | Liesbos | 22042021 | 76720 | 3 | alan | control | |
| 76828 | m | NL | Forest | Liesbos | 22042021 | 76720 | 22 | alan | | died after 1st year |
| 76817 | f | NL | Forest | Oosterhout | 22042021 | 75920 | 35 | control | alan | |
| 35008 | f | NL | Forest | Vlieland | 32042021 | 76656 | 23 | control | alan | |
| 35003 | f | NL | Forest | Vlieland | 32042021 | 76757 | 2 | control | alan | |
| 35005 | m | NL | Forest | Vlieland | 32042021 | 76757 | 7 | alan | control | |
| 35006 | m | NL | Forest | Vlieland | 32042021 | 76757 | 23 | control | alan | |
| 76891 | m | NL | Forest | Vlieland | 32042021 | 76776 | 8 | | alan | replaced BD76755 |
| 76896 | m | NL | Forest | Vlieland | 32042021 | 76776 | 11 | alan | control | |
| 76893 | m | NL | Forest | Vlieland | 32042021 | 76776 | 31 | control | alan | |
| 76890 | f | NL | Forest | Vlieland | 32042021 | 76781 | 7 | alan | control | |
| 76897 | m | NL | Forest | Vlieland | 32042021 | 76796 | 2 | control | alan | |

| | | | | | | | | | | |
|---|---|---|---|---|---|---|---|---|---|---|
| 76900 | m | NL | Forest | Vlieland | 32042021 | 76796 | 12 | | alan | replaced BD76804 |
| 76429 | m | NL | Forest | Wageningen | | Extra | 26 | alan | | died after 1st year |
| 76761 | f | NL | Urban | Amsterdam | 7042021 | 75811 | 1 | alan | control | |
| 76758 | m | NL | Urban | Amsterdam | 7042021 | 75811 | 21 | control | alan | |
| 76760 | m | NL | Urban | Amsterdam | 7042021 | 75821 | 1 | alan | control | |
| 76765 | m | NL | Urban | Amsterdam | 7042021 | 75821 | 4 | control | alan | |
| 76829 | f | NL | Urban | Amsterdam | 14042021 | 75887 | 26 | alan | control | |
| 76786 | f | NL | Urban | Amsterdam | 17042021 | 75892 | 4 | control | alan | |
| 76789 | f | NL | Urban | Amsterdam | 17042021 | 75892 | 21 | control | alan | |
| 76792 | m | NL | Urban | Delft | 13042021 | 75846 | 5 | alan | control | |
| 76755 | m | NL | Urban | Delft | 11042021 | 75847 | 8 | control | | died after 1st year |
| 76838 | f | NL | Urban | Delft | 11042021 | 75849 | 33 | control | alan | |
| 76839 | f | NL | Urban | Delft | 11042021 | 75850 | 8 | control | alan | |
| 76864 | f | NL | Urban | Delft | 28042021 | 76787 | 5 | alan | control | |
| 76754 | m | NL | Urban | Rotterdam | 11042021 | 75849 | 25 | control | alan | |
| 76805 | f | NL | Urban | Rotterdam | 210420201 | 75984 | 12 | control | alan | |
| 76806 | f | NL | Urban | Rotterdam | 210420201 | 75984 | 25 | control | alan | |
| 76825 | m | NL | Urban | Rotterdam | 210420201 | 75984 | 30 | alan | control | |
| 76803 | f | NL | Urban | Rotterdam | 210420201 | 75985 | 9 | alan | control | |
| 76804 | m | NL | Urban | Rotterdam | 210420201 | 75985 | 12 | control | | died after 1st year |
| 76808 | m | NL | Urban | Rotterdam | 210420201 | 76166 | 9 | alan | control | |
| 76807 | f | NL | Urban | Rotterdam | 210420201 | 76166 | 30 | alan | control | |
| 76801 | f | NL | Urban | Utrecht | 12042021 | 75885 | 13 | alan | | died after 1st year |
| 76845 | f | NL | Urban | Utrecht | 20042121 | 76058 | 16 | control | alan | |
| 76846 | m | NL | Urban | Utrecht | 19042021 | 76060 | 13 | alan | | died after 1st year |
| 76847 | f | NL | Urban | Utrecht | 19042021 | 76060 | 29 | control | alan | |
| 76849 | m | NL | Urban | Utrecht | 19042021 | 76060 | 34 | alan | control | |

| | | | | | | | | | |
|---|---|---|---|---|---|---|---|---|---|
| 76856 | m | NL | Urban | Utrecht | 26042021 | 76505 | 29 | control | alan |
| 76853 | m | NL | Urban | Utrecht | 26042021 | 76506 | 16 | control | alan |
| 76857 | f | NL | Urban | Utrecht | 26042021 | 76506 | 34 | alan | control |

**3)** Correlation between feeding activity and activity based on camera observations

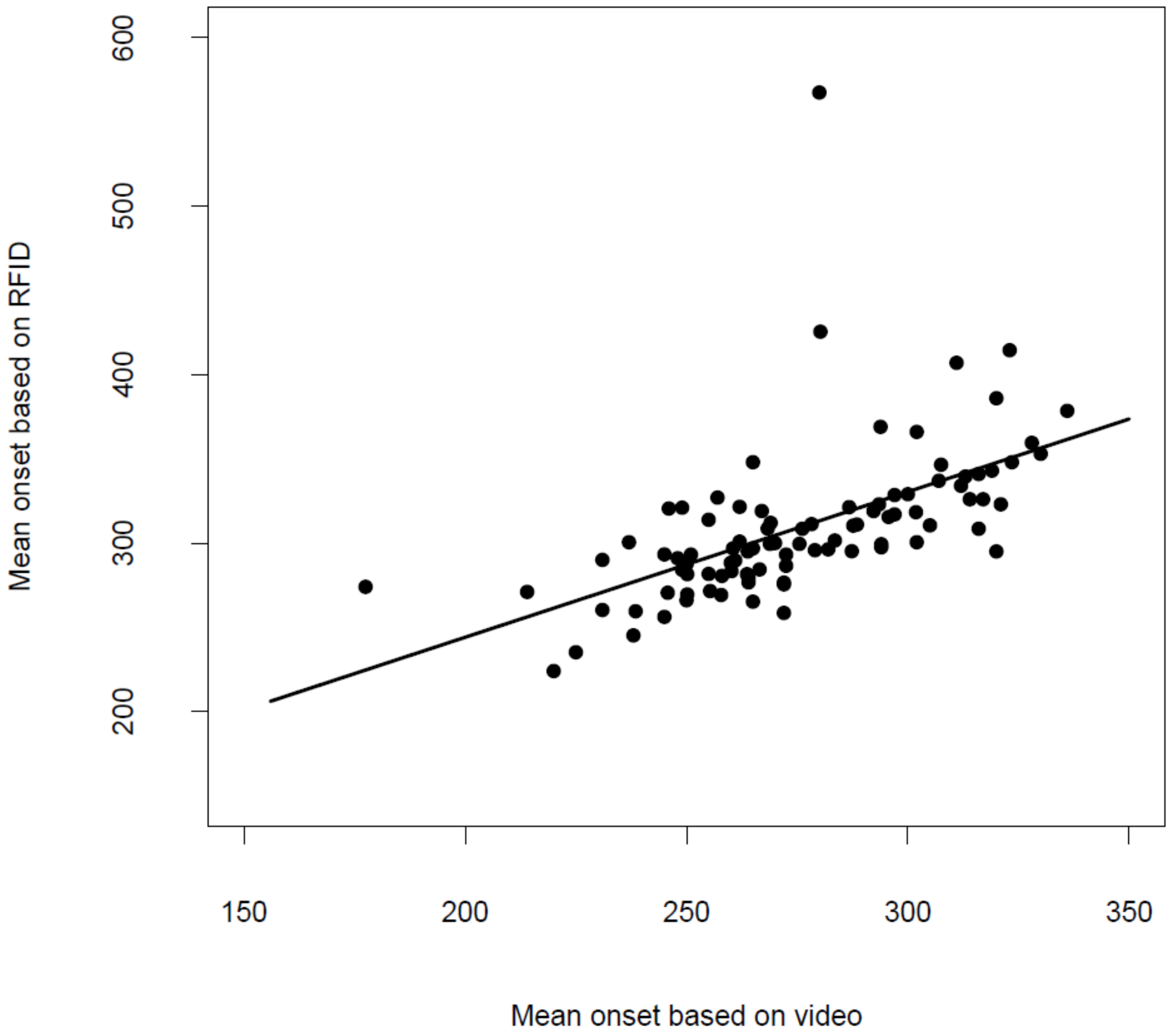

**Figure S3:** Correlation between mean onset of feeding and mean onset of activity based on video recordings in the climate controlled aviaries.

### 4) Spectral composition of lights used in experiments

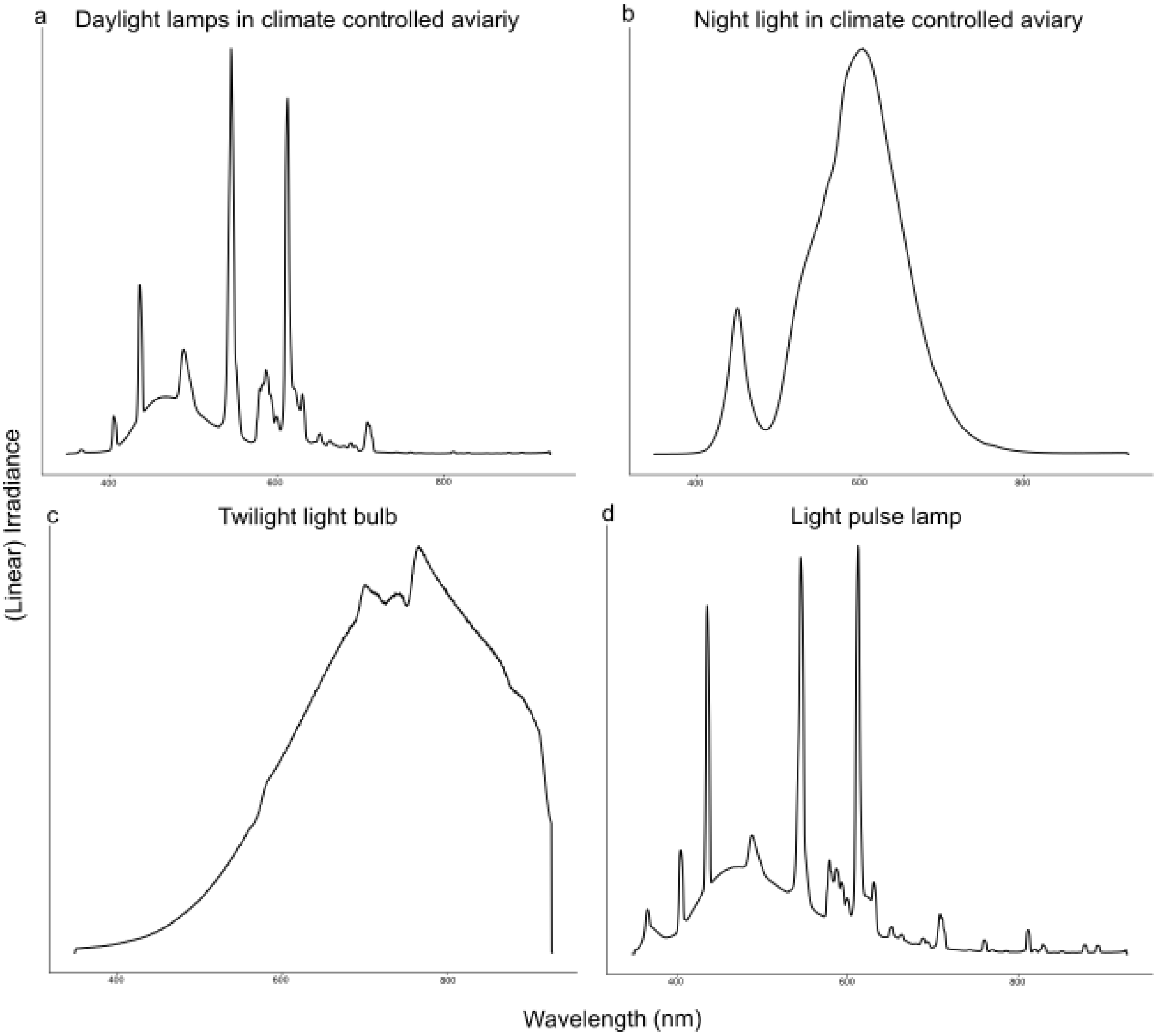

**Figure S4:** Spectral composition of the a) lamps used for daylight in the climate-controlled aviaries, b) night light used for ALAN treatment in the climate-controlled aviaries and as dim light in individual cages for *tau* measurement, c) the light bulb going on 30 min before daylights lamp on and off 30 min after lights off (to mimic twilight phases) and d) the light used for light pulse in individual cages. The measurements were taken using an Ocean Insight OEPro spectrometer with a 200 µm slit and standard cosine corrector.

### 5) Exploring potential populational “driving” effects

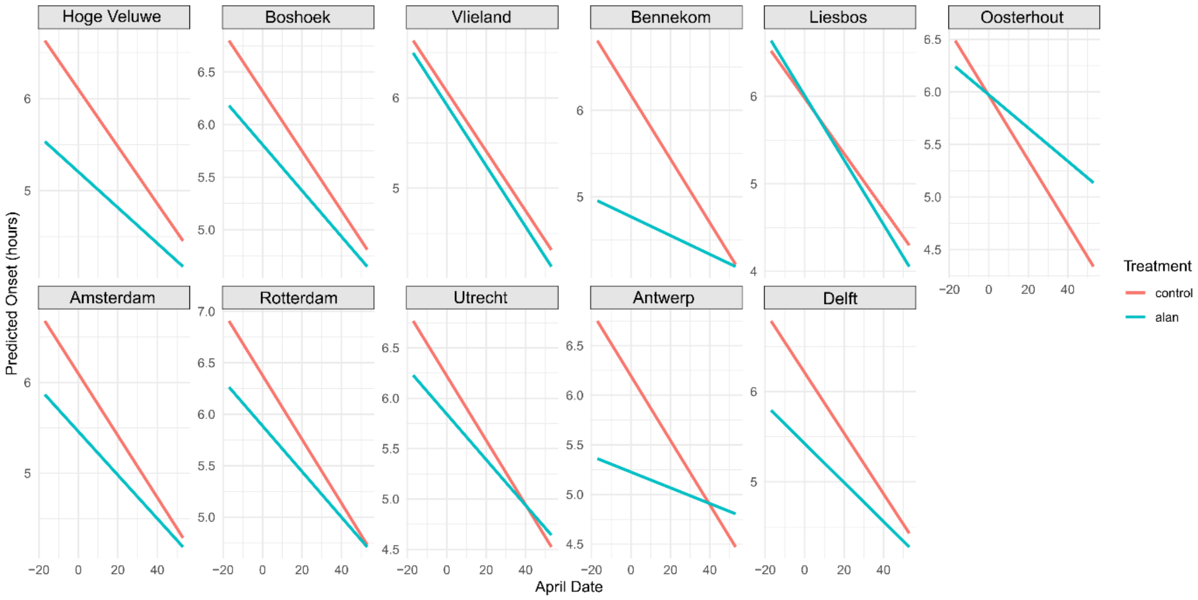

**Figure S5:** Plotted predicted regression lines from model containing Treatment*Area*Aprildate interaction. Regression lines are plotted per population (Forest populations top, city populations below).

6) **Statistical output for activity onset and onset in climate-controlled aviaries**

**Supplementary Table 2: Statistical output for Feeding onset, offset and duration.** Effects of treatment (control or ALAN), Habitat (Forest or city) and April date (which is days in April, proxy for time as day length increase) and the interaction of these variables. The main terms year (2022 or 2023) and sex (male or female) are included as covariates but not tested for. Statistics are given at the point of exclusion of the model, using Kenward Roger approximation through the function "KRmodcomp". For feeding onset full model was the final model, while for feeding offset final model consisted of the interaction between treatment and April date and Habitat and April date. Coefficients reported are for the final model with intercept removed facilitate interpretation and random effects.

| **Feeding onset** | **Estimate** | **SE** | **Ndf** | **ddf** | ***F-value*** | ***p-value*** |
|---|---|---|---|---|---|---|
| Treatment:Habitat:AprilDate | | | 1.00 | 6494.11 | 5.21 | **0.022** |
| Treatment (control) | 6.3316 | 0.0826 | | | | |
| Treatment (ALAN) | 5.6901 | 0.0828 | | | | |
| Habitat (city) | -0.0009 | 0.1051 | | | | |
| Aprildate | -0.0327 | 0.0009 | | | | |
| Year (2023) | 0.0305 | 0.0191 | | | | |
| Sex (male) | -0.3378 | 0.0646 | | | | |
| Treatment(ALAN):Habitat(city) | -0.001 | 0.05 | | | | |
| Treatment(ALAN):AprilDate | 0.008 | 0.001 | | | | |
| Habitat(city):AprilDate | 0.0004 | 0.0013 | | | | |
| Treatment(alan):Habitat(city):AprilDate | 0.0044 | 0.0019 | | | | |
| **Random effects** | **Variance** | **SD** | | | | |
| Individual ID:Aviary | 0.05 | 0.2246 | | | | |
| Aviary | 0.0217 | 0.1475 | | | | |
| Mother ID:population | 0.0355 | 0.1885 | | | | |
| Population | 0.0032 | 0.0561 | | | | |
| Residual | 0.4205 | 0.6485 | | | | |
| *Number of obs: 6528, groups: Individual ID:Aviary, 81; Aviary, 42; Mother:population, 41; population, 11* | | | | | | |
| **Feeding Offset** | **Estimate** | **SE** | **Ndf** | **ddf** | ***F-value*** | ***p-value*** |
| Treatment:Habitat:AprilDate | | | 1.00 | 6475.78 | 2.12 | 0.14 |
| Treatment:Habitat | | | 1.00 | 6321.31 | 0.52 | 0.47 |
| Treatment:Aprildate | | | 1.00 | 6429.13 | 11.162 | **0.0008** |
| Habitat:Aprildate | | | 1.00 | 6488.7 | 80.09 | **< 0.0001 ***** |
| Treatment (control) | 18.6536 | 0.0557 | | | | |
| Treatment (ALAN) | 18.7104 | 0.0557 | | | | |
| Habitat (city) | 0.0238 | 0.0658 | | | | |
| Aprildate | 0.0314 | 0.0007 | | | | |
| Year (2023) | 0.2955 | 0.0166 | | | | |
| Sex (male) | 0.2022 | 0.0457 | | | | |
| Treatment(ALAN):AprilDate | -0.0028 | 0.0008 | | | | |
| Habitat(city):AprilDate | -0.0070 | 0.0007 | | | | |
| **Random effects**: | **Variance** | **SD** | | | | |
| Individual ID:Aviary | 0.0284 | 0.1684 | | | | |
| Aviary | 0.0174 | 0.1318 | | | | |
| Mother ID:population | 0.0086 | 0.0927 | | | | |
| Population | 0.0 | 0.0 | | | | |
| Residual | 0.3139 | 0.5602 | | | | |
| *Number of obs: 6524, groups: Individual ID:Aviary, 81; Aviary, 42; Mother:population, 41; Population, 11* | | | | | | |

7) **Statistical output for the three clock properties measures (first onset, *tau and response to light pulse***

**Supplementary Table 3: statistical output for the clock properties: First onset in constant conditions (clock timing), *Tau* (Clock speed) and phase shift (Clock response to light pulse; CT0).** Effects of treatment (dark control or ALAN) and habitat (city or forest) and the interaction of these variables. The main terms year (2022 or 2023), sex (male or female) and group (A or B) are included as covariates but not tested for. Statistics are given at the point of exclusion of the model, using Kenward Roger approximation through the function “KRmodcomp”.

| **First onset** | **Estimate** | **SE** | **Ndf** | **ddf** | ***F-value*** | ***p-value*** |
|---|---|---|---|---|---|---|
| Treatment:Habitat | | | 1.00 | 56.3035 | 0.4791 | 0.49 |
| Treatment | | | 1.00 | 58.3347 | 0.5454 | 0.46 |
| Habitat | | | 1.00 | 7.2808 | 0.0001 | 0.99 |
| Group A | -0.4396 | 0.1313 | | | | |
| Group B | -0.0433 | 0.1326 | | | | |
| Year (2023) | -0.1110 | 0.1586 | | | | |
| Sex (male) | -0.3229 | 0.135 | | | | |
| ***Tau*** | | | | | | |
| Treatment:Habitat | | | 1.00 | 61.7822 | 0.0002 | 0.9887 |
| Treatment | | | 1.00 | 62.9053 | 0.3325 | 0.5662 |
| Habitat | | | 1.00 | 7.7752 | 0.2281 | 0.6461 |
| Group A | 23.4058 | 0.0440 | | | | |
| Group B | 23.3417 | 0.0445 | | | | |
| Year (2023) | 0.0242 | 0.0298 | | | | |
| Sex (male) | -0.0606 | 0.0479 | | | | |
| **Phase shift** | | | | | | |
| Treatment:Habitat | | | 1.00 | 53.1648 | 0.4644 | 0.4985 |
| Treatment | | | 1.00 | 54.0984 | 0.1022 | 0.7505 |
| Habitat | | | 1.00 | 7.3361 | 0.1788 | 0.6845 |
| Group A | 1.8604 | 0.1752 | | | | |
| Group B | 1.4895 | 0.1615 | | | | |
| Year (2023) | -0.4498 | 0.1349 | | | | |
| Sex (male) | -0.2770 | 0.1703 | | | | |

### 8) Individual slopes for ALAN and control conditions in climate controlled aviaries

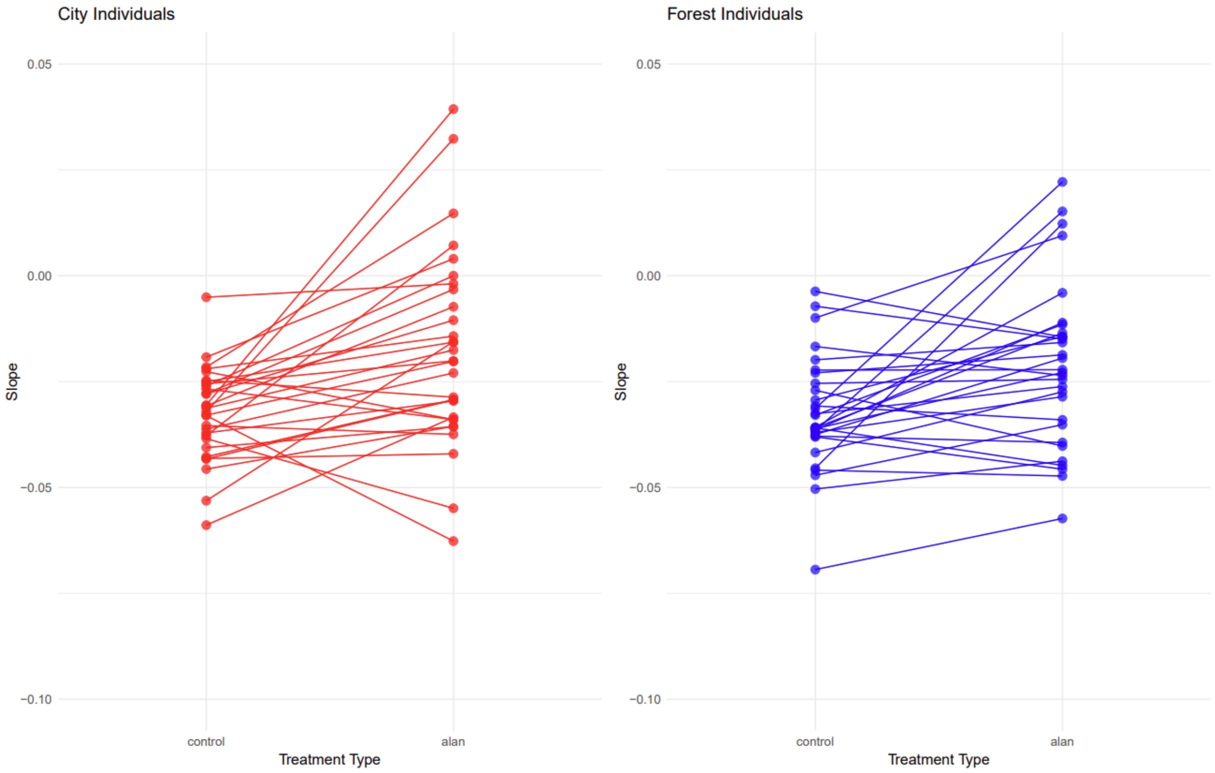

**Figure S5:** Individual slope for activity onset for birds under control and ALAN treatment in the climate controlled aviaries. Only birds present for both years (2022 and 2023) used. To calculate individual slope a linear model was fitted with onset as response and individual ID : Aprildate interaction. The difference between the slope under control vs ALAN treatment was calculated (Δ onset slope = Individual control – Individual ALAN).
